# The genetic overlap between mood disorders and cardio-metabolic diseases: A systematic review of genome wide and candidate gene studies

**DOI:** 10.1101/150615

**Authors:** Azmeraw T. Amare, Klaus Oliver Schubert, Sarah Cohen-Woods, Bernhard T. Baune

## Abstract

Meta-analyses of genome-wide association studies (meta-GWAS) and candidate gene studies have identified genetic variants associated with cardiovascular diseases, metabolic diseases, and mood disorders. Although previous efforts were successful for individual disease conditions (single disease), limited information exists on shared genetic risk between these disorders. This article presents a detailed review and analysis of cardio-metabolic diseases risk (CMD-R) genes that are also associated with mood disorders. Firstly, we reviewed meta-GWA studies published until January 2016, for the diseases “type 2 diabetes, coronary artery disease, hypertension” and/or for the risk factors “blood pressure, obesity, plasma lipid levels, insulin and glucose related traits”. We then searched the literature for published associations of these CMD-R genes with mood disorders. We considered studies that reported a significant association of at least one of the CMD-R genes and “depressive disorder” OR “depressive symptoms” OR “bipolar disorder” OR “lithium treatment”, OR “serotonin reuptake inhibitors treatment”. Our review revealed 24 potential pleiotropic genes that are likely to be shared between mood disorders and CMD-Rs. These genes include *MTHFR*, *CACNA1D*, *CACNB2*, *GNAS*, *ADRB1*, *NCAN*, *REST*, *FTO*, *POMC*, *BDNF*, *CREB*, *ITIH4*, *LEP*, *GSK3B*, *SLC18A1*, *TLR4*, *PPP1R1B*, *APOE*, *CRY2*, *HTR1A*, *ADRA2A*, *TCF7L2*, *MTNR1B*, and *IGF1*. A pathway analysis of these genes revealed significant pathways: corticotrophin-releasing hormone signaling, AMPK signaling, cAMP-mediated or G-protein coupled receptor signaling, axonal guidance signaling, serotonin and dopamine receptors signaling, dopamine-DARPP32 feedback in cAMP signaling, circadian rhythm signaling and leptin signaling. Our findings provide insights in to the shared biological mechanisms of mood disorders and cardio-metabolic diseases.

## INTRODUCTION

Major depressive disorder (MDD), bipolar disorder (BPD), coronary artery diseases, type 2 diabetes and hypertension are amongst the major causes of disability, morbidity and mortality worldwide (1, 2). While each of these conditions independently represent a major burden facing the health-care systems (1-3), their co-occurrence (co-morbidity) aggravates the situation and represents a challenge in psychosomatic medicine. Epidemiologically, MDD and BPD are bi-directionally associated with cardio-metabolic diseases (4, 5). One explanation for these relationships could be the presence of pleiotropic (common) genes and shared biological pathways that function as a hub encoding for proteins connecting the disorders. Potential common biological mechanisms underlying mood disorders and cardio-metabolic disease comorbidity have been proposed, including altered circadian rhythms (6), abnormal hypothalamic-pituitary-adrenal axis (HPA axis) function (7), imbalanced neurotransmitters (8), and inflammation (5). However, the molecular drivers of these commonly affected mechanisms remain poorly understood.

### The genetics of mood disorders and cardio-metabolic diseases

Major depression, bipolar disorder and cardio-metabolic diseases are highly heritable and caused by a combination of genetic and environmental factors. Genetic factors contribute to 31-42% in MDD (9), 59% - 85% in BPD (10, 11), 30-60% in coronary artery diseases (12), 26-69% in type 2 diabetes (13, 14), 24-37% in blood pressure (15), 40–70% in obesity (16), and 58-66% in serum lipids level (17). Moreover, twin studies have revealed high genetic co-heritabilities (genetic correlations) between mood disorders and the different cardio-metabolic disorders suggesting the influence of pleiotropic genes and shared biological pathways among them. For instance, the genetic correlation of depression with hypertension is estimated to be 19%, and between depression and heart disease is about 42% (18). The genetic correlation of depressive symptoms with plasma lipids level ranges from 10% to 31% (19), and 12% of the genetic component for depression is shared with obesity (20).

In the last decade, substantial amounts of univariate (single disease) meta-GWA studies and candidate gene studies have been published. Indeed, the meta-GWA studies and candidate gene studies have successfully identified a considerable list of candidate genes for major depressive disorder (21, 22), bipolar disorder (23), coronary artery diseases (24), type 2 diabetes (25), hypertension (26), obesity (27), plasma lipids level (28), insulin and glucose traits (25, 29, 30), and blood pressure (26, 31).

Despite the potential significance of studying pleiotropic genes and shared biological pathways, previous meta-GWAS and candidate gene studies were entirely focused on a single phenotype approach (single disease). A recent analysis of SNPs and genes from the NHGRI GWAS catalogue (32) has showed as 16.9% of the genes and 4.6% of the SNPs have pleiotropic effects on complex diseases (33). Considering such evidence, we hypothesized that common genetic signatures and biological pathways mediate the mood disorders to cardio-metabolic diseases relationship. Additionally, these genes and their signalling pathways can influence the response to treatments in mood disorder patients (figure 1). In this review, we systematically investigated the CMD-R genes that are possibly associated with mood disorders susceptibility, and with treatment response to MDD and BPD. We performed pathway and gene network analyses of these genes to provide additional insights in to the common pathways and biological mechanisms regulating mood disorders and the CMD-Rs. Understanding of these common pathways may provide new insights and novel ways for the diagnosis and treatment of comorbid cardio-metabolic and mood disorders.

**Figure 1.**
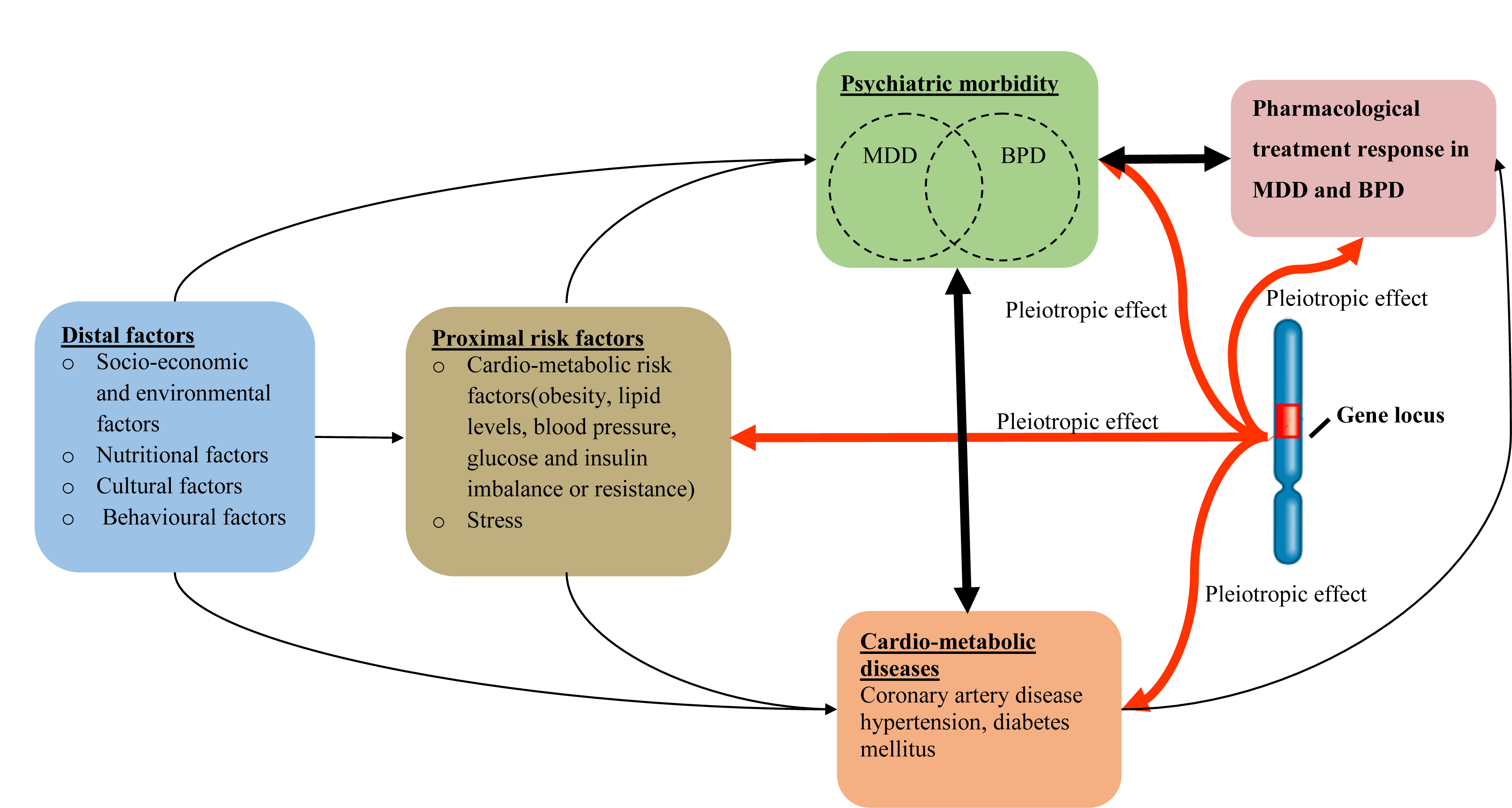
Schematic model of the potential pleiotropic effects of a shared gene locus that is associated with mood disorders and cardio-metabolic diseases. The red bold lines represent the pleiotropic effect of a genetic locus on CMD-Rs, MDD, BPD and treatment response in MDD and BPD. The bidirectional arrows indicate bidirectional relationships. MDD: Major depressive disorder, BPD: Bipolar disorder, CMD-Rs: cardio-metabolic diseases and risk factors.

## METHODS AND MATERIALS

### SEARCH STRATEGY

#### Step 1: Identification of published candidate genes for cardio-metabolic diseases

We carried out a systematic search of candidate genes for the cardio-metabolic diseases and/or associated risk factors. The National Human Genome Research Institute (NHGRI) GWAS catalogue (32), Westra et al., 2013 (34) and Multiple Tissue Human Expression Resource (MuTHER)(35) databases were used to identify the CMD-R genes. We reviewed meta-GWA study papers published until January 2016 for the diseases “type 2 diabetes” OR “coronary artery disease” OR “hypertension” and (or) for the risk factors “blood pressure”OR “obesity or body mass index (BMI)” OR “plasma lipid levels (high-density lipoprotein, low-density lipoprotein, triglycerides, total cholesterol)” OR “insulin and glucose related traits (fasting glucose, fasting insulin, fasting proinsulin, insulin resistance-HOMA-IR), beta cell function-HOMA-β and glycated haemoglobinA1C-HbA1C)”.

GWAS significant SNPs information (lead SNPs, reported genes, author (s), PubMed ID, date of publication, journal, discovery and replication sample sizes) was downloaded from the GWAS catalogue database. Additional information about the effect of the lead SNPs on nearby gene expression (cis-eQTLs) was collected from their respective publications. For the SNPs with no cis-eQTL information in their respective publications, we performed expression quantitative trait loci (cis-eQTL) analysis to verify the functional relationship between the reported genes and the lead SNPs using two publicly available databases: Westra et al., 2013 (34), and MuTHER (35). A CMD-R gene was considered as a candidate gene if, 1) it was nearby to the lead SNP and its expression was influenced by the lead SNP (cis-eQTL); or 2) the genes were biologically well known to influence at least one of the CMD-Rs. We took the identified CMD-R genes forward for the second literature review, as described below.

#### Step 2: Exploration of the role of cardio-metabolic genes in mood disorders

In the second systematic review, we conducted a literature search in PubMed (MEDLINE database) for any genome wide association, candidate gene, or gene expression analysis study published in the fields of mood disorders and mood disorders pharmacogenetics until January 2016. We considered studies that reported at least one of the CMD-R genes for “depressive disorder” OR “depressive symptoms” OR “bipolar disorder” OR “mood disorder” OR “lithium treatment”, OR “Selective Serotonin Reuptake Inhibitors (SSRIs) treatment”.

### BIOLOGICAL PATHWAY AND NETWORK ANALYSIS

The potential pleiotropic genes were further explored to identify the most enriched canonical pathways and visualize gene networks using QIAGEN’s Ingenuity^®^ Pathway Analysis (IPA^®^, QIAGEN Redwood City, www.qiagen.com/ingenuity). For the analysis, all the twenty-four potential pleiotropic genes were entered into the software. IPA compares the proportion of input genes mapping to a biological pathway to the reference genes in the Ingenuity databases. The significance of the overrepresented canonical pathways were determined using the right-tailed Fisher”s exact test. After correction for multiple testing, significance levels were expressed as the IPA p-value. A gene networks that connects the input genes with MDD, BPD and the cardio-metabolic disorders was also generated.

## RESULTS

### Characteristics of meta-GWA studies for the cardio-metabolic disorders

The literature search in the GWAS catalogue yielded 153 meta-GWA studies for the CMD-Rs: 38 studies for type 2 diabetes, 17 studies for coronary artery disease, 15 studies for hypertension and blood pressure, 26 studies for obesity (BMI), 37 studies for lipids and 20 studies for glucose and insulin traits (figure 2). As shown in figure 2, the meta-GWA studies reported 1047 lead SNPs and 682 nearby genes. Of these, 123 genes were biologically relevant to the cardio-metabolic diseases and associated risk factors. These genes were reviewed for their association with mood disorders and pharmacogenetics of mood disorders. Twenty-four of the 123 genes have been implicated in mood disorders and/or pharmacogenetics of mood disorders; and we named these genes the Cardio-Metabolic Mood disorders hub (CMMDh) genes.

**Figure 2:**
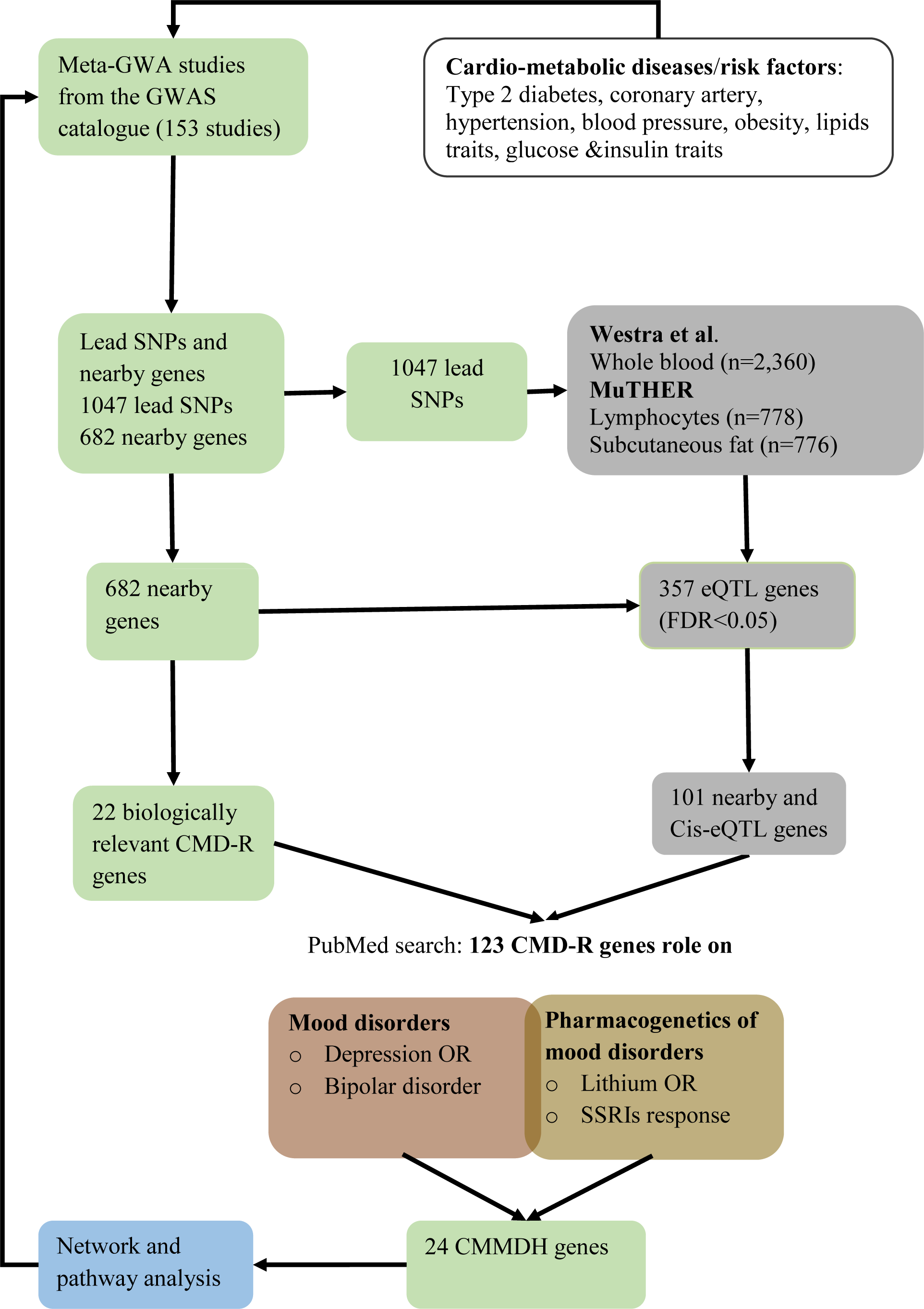
The flow chart shows the stages of literature search and evaluation of candidate pleiotropic genes for the CMD-Rs and mood disorders. CMD-R genes refers to the genes that were biologically well-known to influence at least one of the CMD-Rs or those genes nearby to the lead SNPs and their expression was influenced by the lead SNPs (cis-eQTL). CMD-R: Cardio-metabolic diseases and associated risk factors, MuTHER: Multiple Tissue Human Expression Resource, CMMDh: Cardio-Metabolic Mood Disorders hub genes, Cis-eQTL: Cis (nearby) gene expression quantitative trait loci.

**Figure 2:**
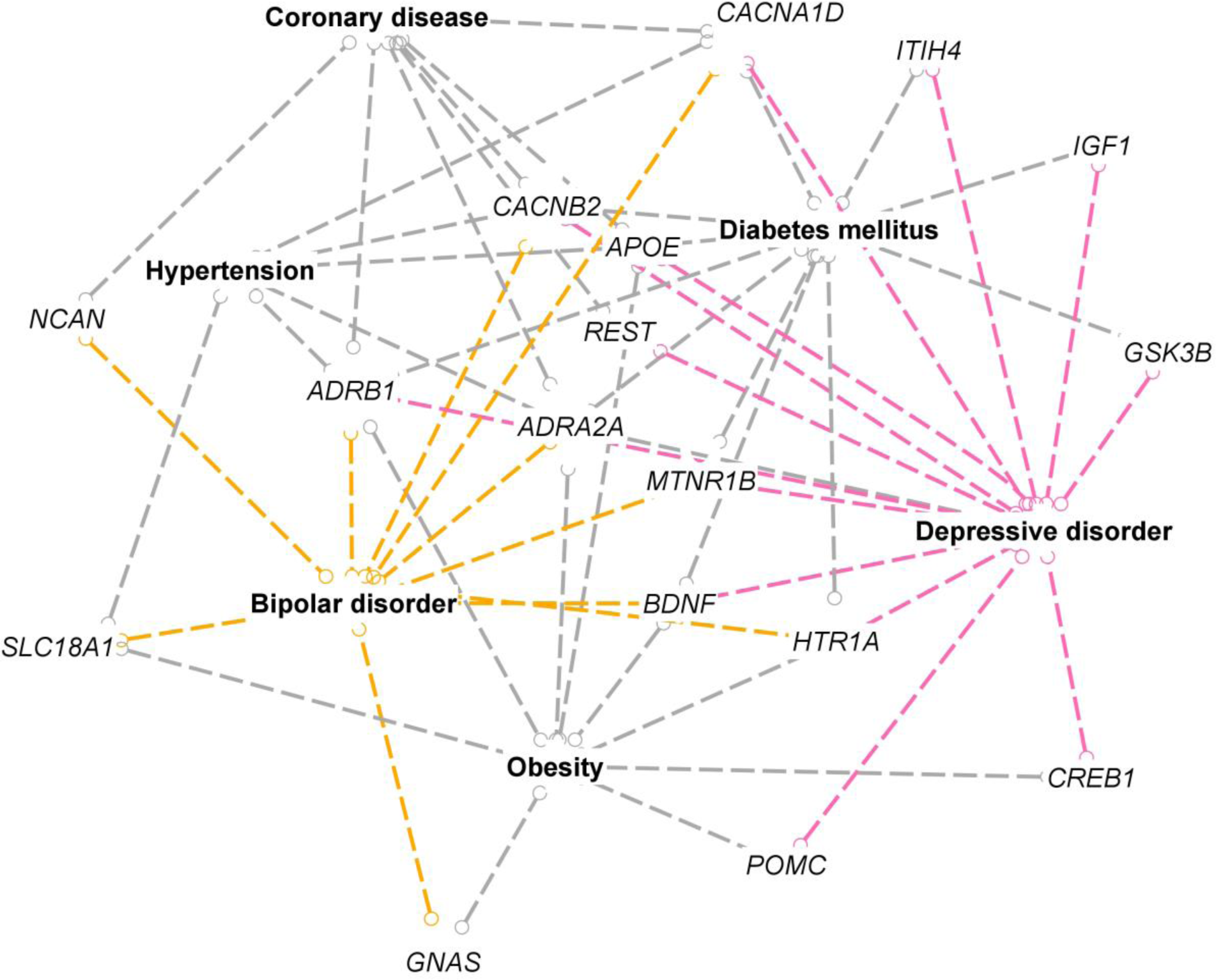
IPA-generated network of genes, as indicated by the dashed lines, shared between coronary artery diseases, hypertension, diabetes mellitus, obesity, MDD and BPD, highlighting CMMDh genes that were related to bipolar disorder (orange) and depression (red).

Table-1 summarizes the 24 CMMDh genes and specific genetic variants across mood disorders and cardio-metabolic diseases. These genes are *MTHFR*, *CACNA1D*, *CACNB2*, *GNAS*, *ADRB1*, *NCAN*, *REST*, *FTO*, *POMC*, *BDNF*, *CREB*, *ITIH4*, *LEP*, *GSK3B*, *SLC18A1*, *TLR4*, *PPP1R1B*, *APOE*, *CRY2*, *HTR1A*, *ADRA2A*, *TCF7L2*, *MTNR1B*, and *IGF1* (for further details see table 1). These genes were over-represented in the following biological pathways; corticotrophin-releasing hormone signaling (*BDNF*, *CREB1*, *GNAS*, *POMC)*; AMPK signaling (*ADRA2A*, *ADRB1*, *CREB1*,*GNAS*, *LEP)*; cAMP-mediated and G-protein coupled receptor signaling *(ADRA2A*, *ADRB1*, *CREB1*, *GNAS*, *HTR1A)*; axonal guidance signaling (*BDNF*, *GNAS*, *GSK3B*, *IGF1*); serotonin and dopamine receptors signaling *(GNAS*, *HTR1A*, *SLC18A1, PPP1R1B)*; dopamine-DARPP32 feedback in cAMP *(PPP1R1B*, *CACNA1D*, *CREB1*, *GNAS)*; leptin signaling (*GNAS*, *LEP*, *POMC)* and the circadian rhythm signaling(*CRY2*,*CREB1*) (table 2).

**Table 1:**
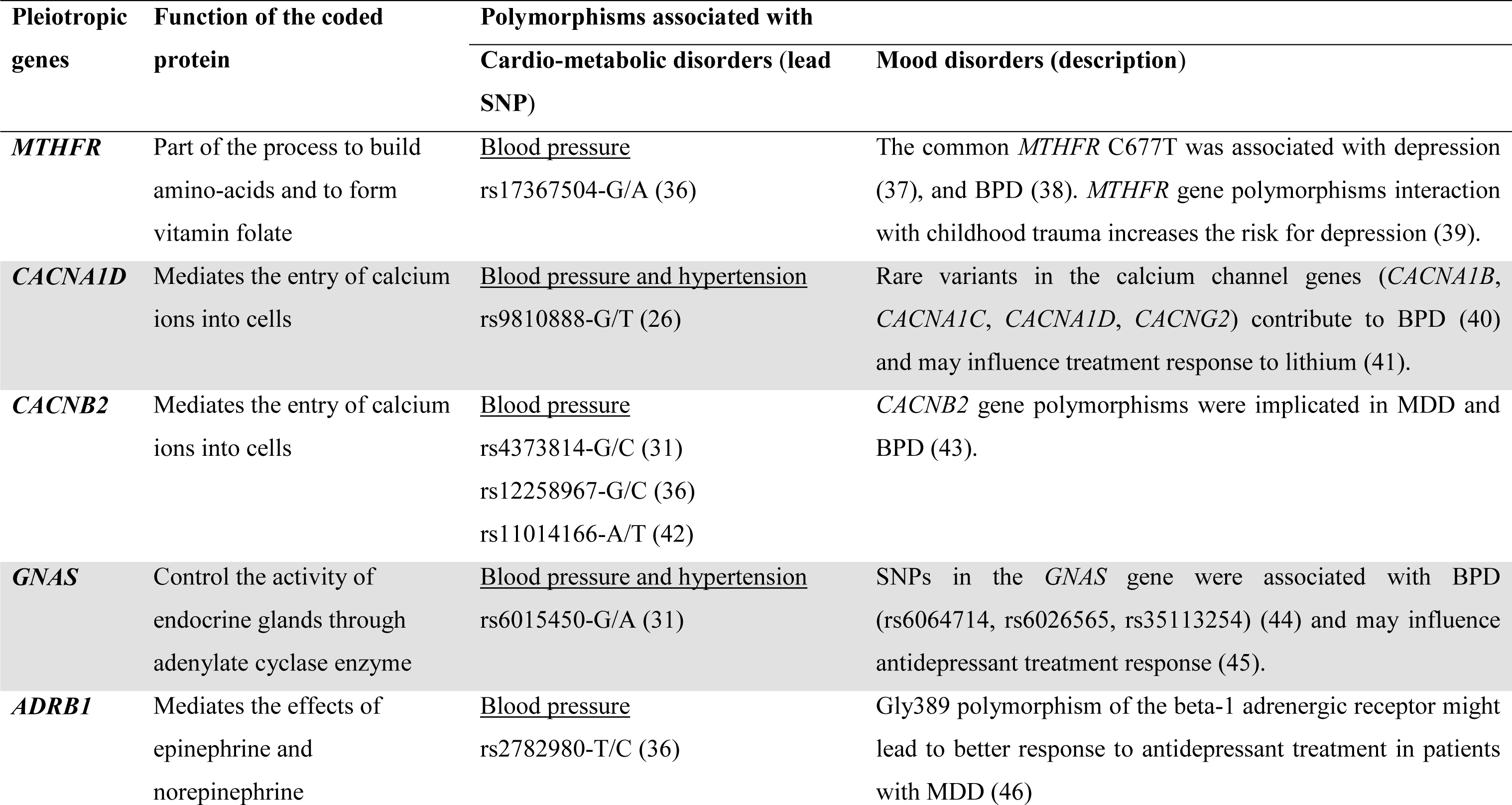
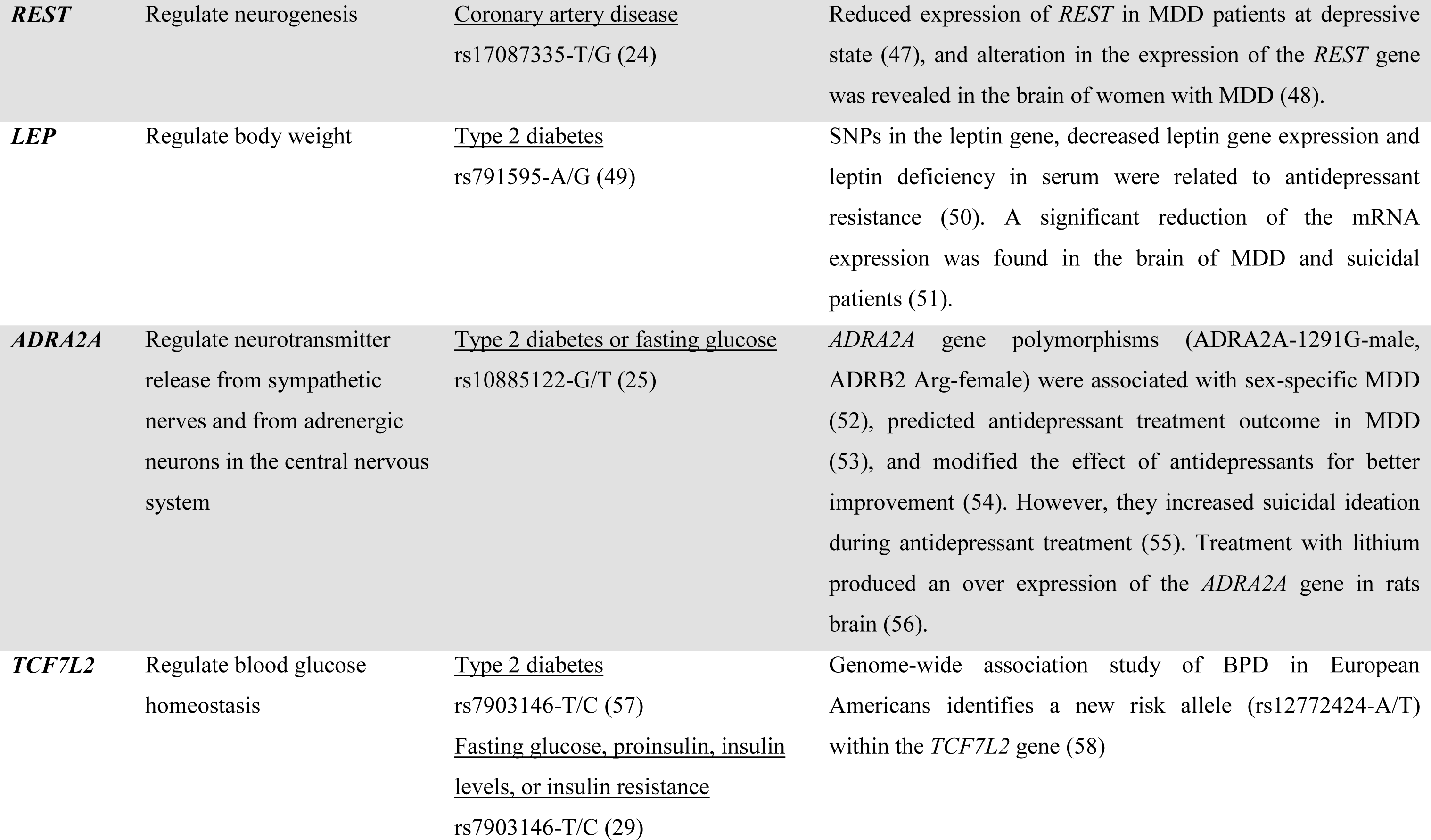
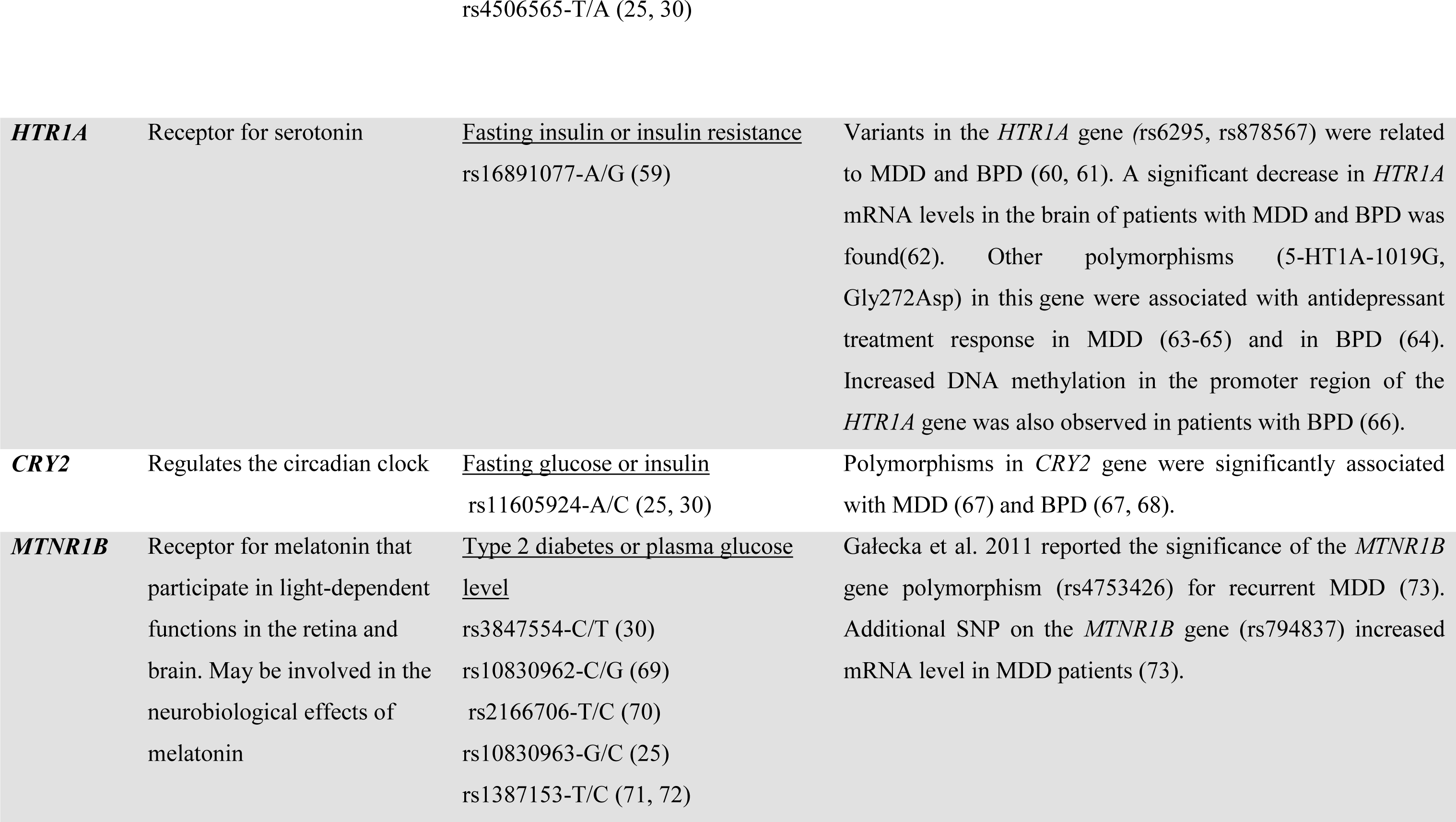
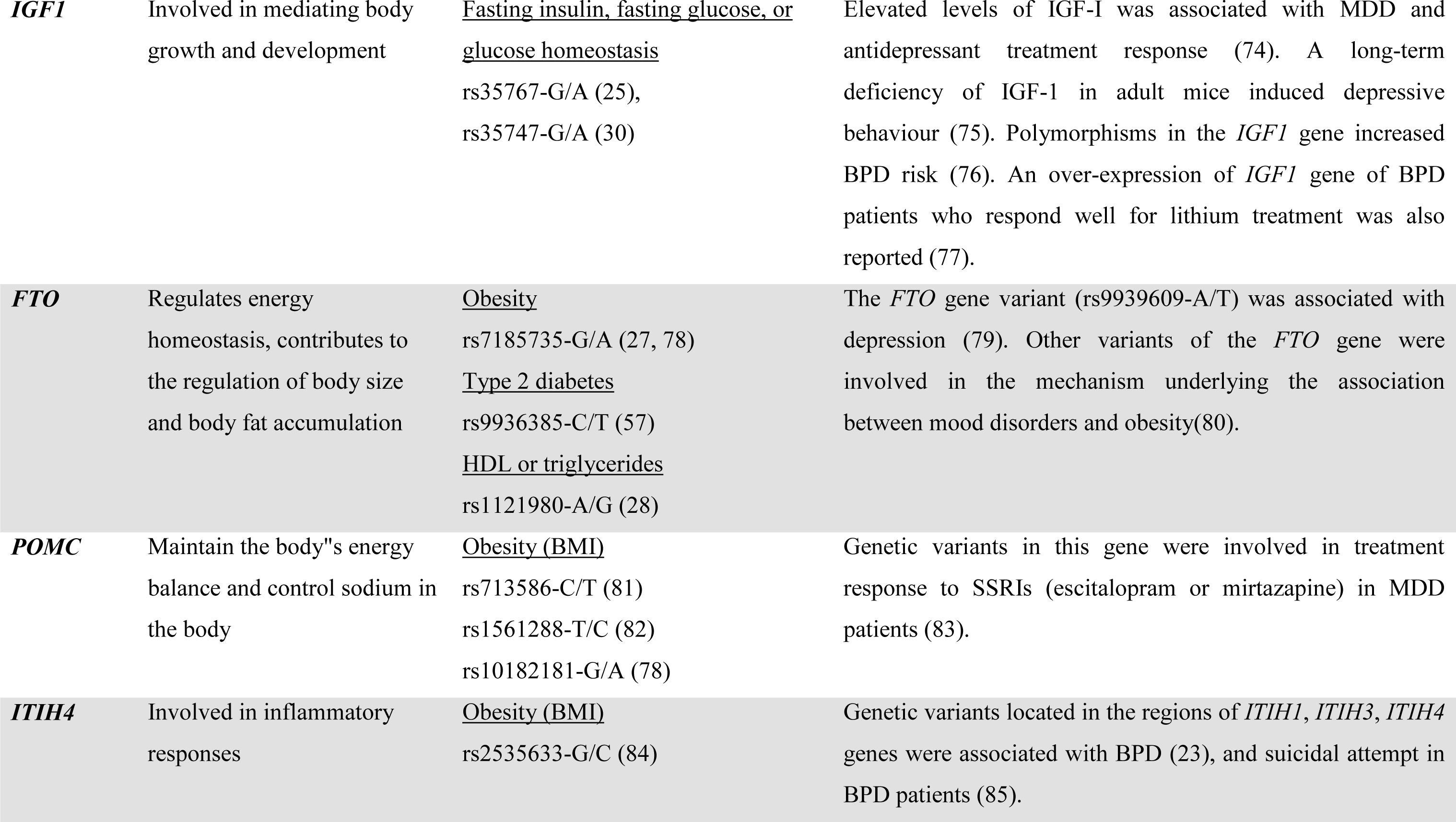
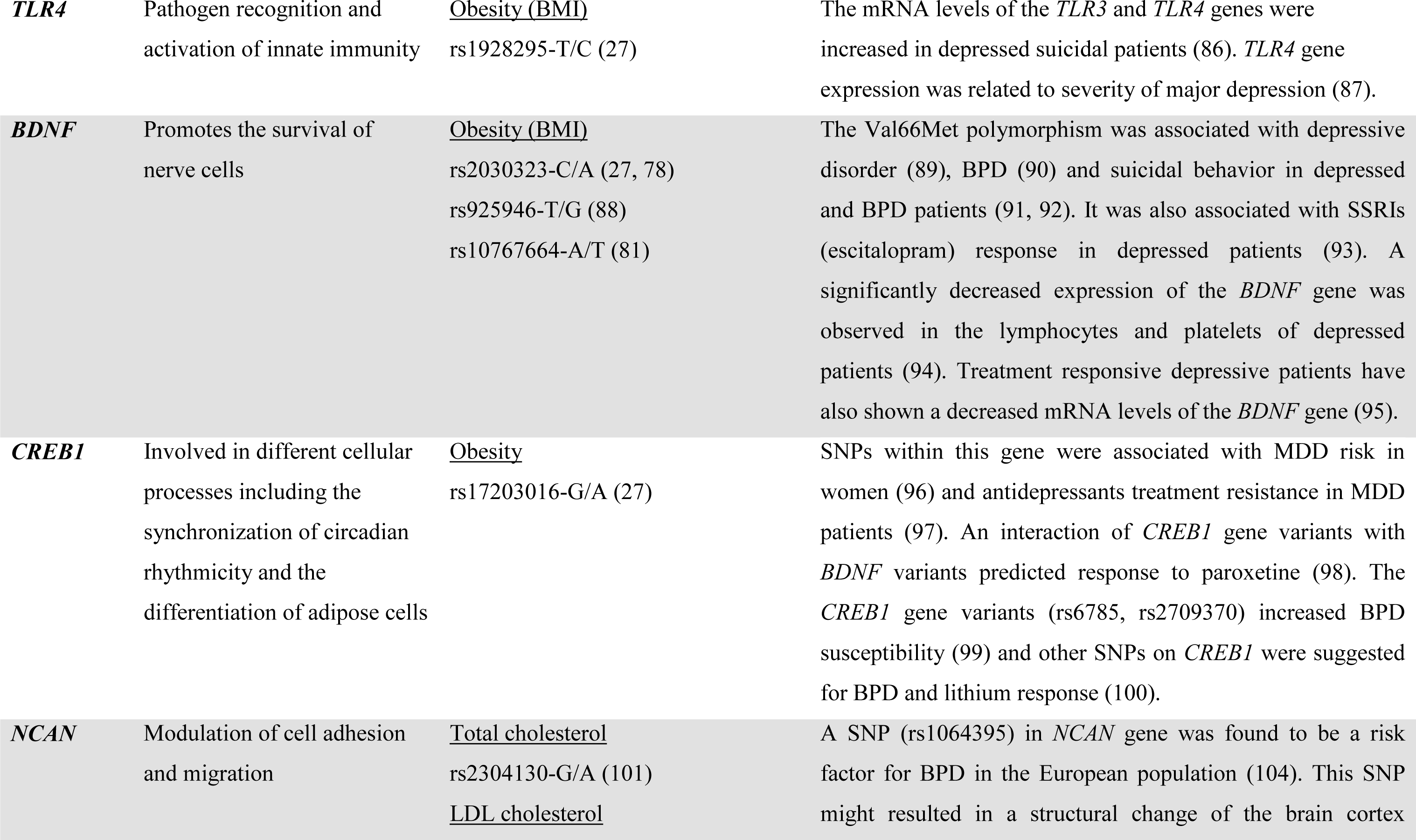
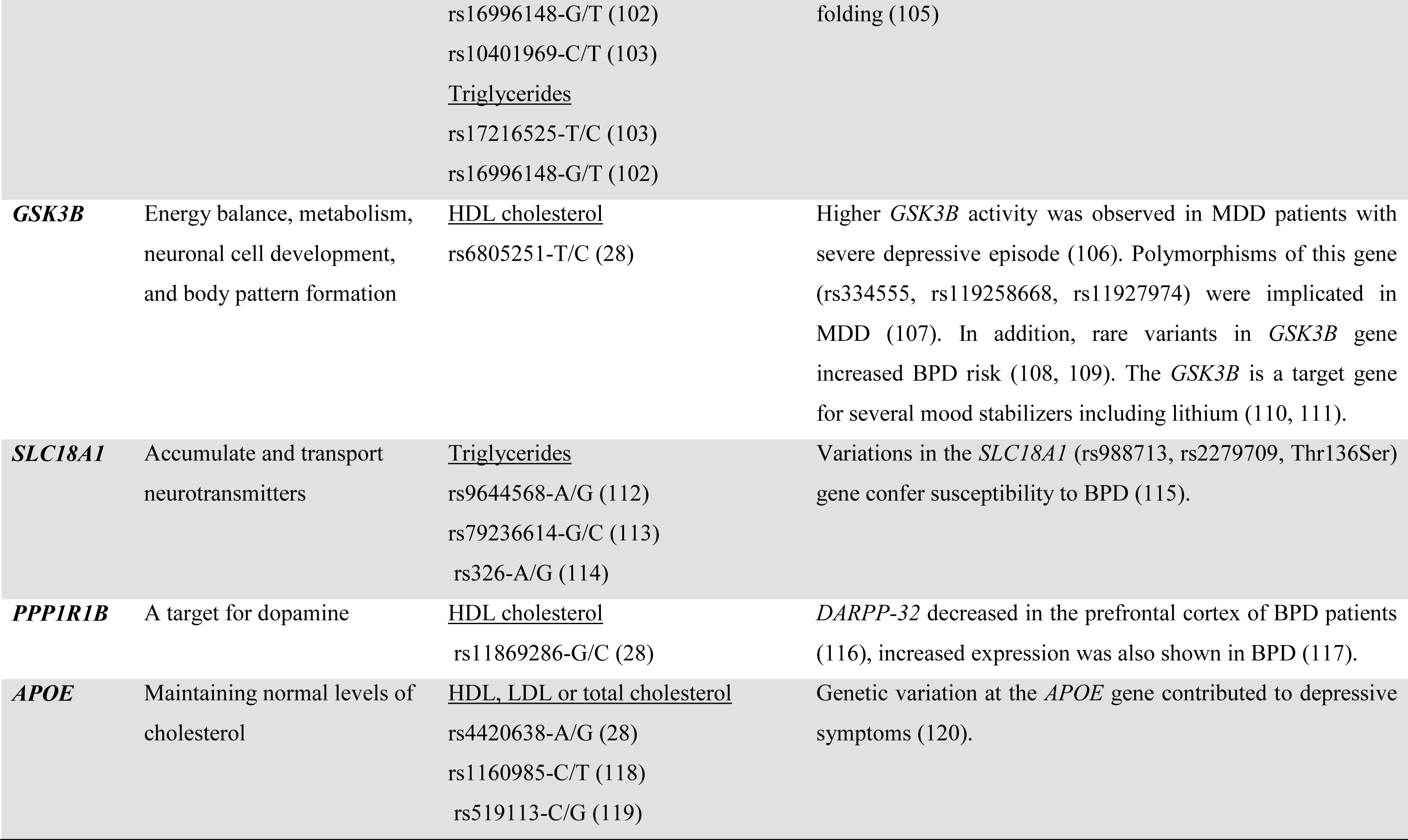
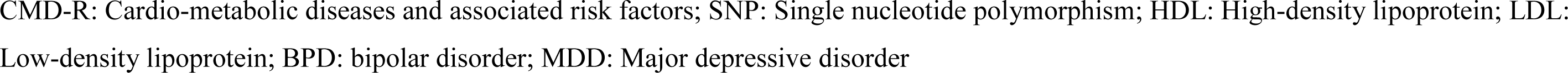
An overview of the 24 CMMDh genes shared between mood disorders and the cardio-metabolic diseases

**Table 2:** The top canonical signaling pathways enriched for the cardio-metabolic mood disorders hub genes

| Canonical pathways | Enriched genes | P-value |
| --- | --- | --- |
| Corticotrophin releasing hormone | *BDNF, CREB1, GNAS, POMC* | $7.77 \times 10^{-6}$ |
| AMPK signaling | *ADRA2A, ADRB1, CREB1, GNAS, LEP* | $1.68 \times 10^{-6}$ |
| cAMP-mediated | *ADRA2A, ADRB1, CREB1, GNAS, HTR1A* | $4.65 \times 10^{-6}$ |
| G-Protein coupled receptor | | $9.92 \times 10^{-6}$ |
| Dopamine-DARPP32 feedback in cAMP | *CACNA1D, CREB1, GNAS, PPP1R1B* | $3.36 \times 10^{-5}$ |
| Serotonin receptor | *GNAS, HTR1A, SLC18A1* | $1.78 \times 10^{-5}$ |
| Dopamine receptor | *SLC18A1, GNAS, PPP1R1B* | $9.94 \times 10^{-5}$ |
| Axonal guidance | *BDNF, GNAS, GSK3B, IGF1* | $1.47 \times 10^{-3}$ |
| Leptin signaling | *GNAS, LEP, POMC* | $8.5 \times 10^{-5}$ |
| Cardiac hypertrophy | *ADRA2A, ADRB1, CACNA1D, CREB1, GNAS, GSK3B, IGF1* | $4.65 \times 10^{-9}$ |
| Circadian rhythm signaling | *CRY2, CREB1* | $6.7 \times 10^{-4}$ |

The table shows the top canonical pathways and enriched CMMDh genes as identified by IPA (P<0.05). The P-value indicates the likelihood of finding gene enrichment of the given pathway by chance.

AMPK: 5’ adenosine monophosphate-activated protein kinase, cAMP: Cyclic Adenosine 3’,5’-monophosphate, CMMDh: Cardio-Metabolic Mood Disorders hub genes.

## DISCUSSION

This is the first cross-disorder review that systematically evaluated candidate pleiotropic genes and biological pathways that are likely to be shared with mood disorders, cardiovascular diseases and metabolic disorders. We revealed 24 cardiovascular and metabolic disease genes implicated in either depression, bipolar disorder or both. These genes belong to interrelated signaling pathways important in the hypotheses of both cardio-metabolic diseases and mood disorders: corticotrophin-releasing hormone signaling, AMPK signaling, cAMP-mediated and G-protein coupled receptor signaling, axonal guidance signaling, serotonin and dopamine receptors signaling, dopamine-DARPP32 Feedback in cAMP signaling, leptin signaling and circadian rhythm signaling.

The corticotrophin-releasing hormone (CRH) signaling is one of the top canonical pathways that may underlie the link between CMD-Rs and mood disorders. The CRH signaling pathway comprises of CRH, CRH receptors (CRHR1, CRHR2), and other CRH-related peptides. It is the principal regulator of the hypothalamic–pituitary–adrenal (HPA) axis. There are consistent findings in the literature that support the role of the HPA axis dysregulation in mediating the risk of mood disorders and cardiovascular outcome (121). Our analysis found enriched CMMDh genes in the CRH signaling pathways (*BDNF*, *CREB1*, *GNAS*, *POMC*). Genetic variants of the genes for BDNF, CREB1, GNAS, and POMC increased the risk of MDD (89, 96), BPD (44), obesity (27, 81), blood pressure and hypertension (31, 36). These genes could be stress responsive, and their activity could be modulated through the activation of the HPA-axis. In animal studies, the expression of *BDNF* (122) and *CREB1* (123) genes was dysregulated by chronic stress. It is therefore possible that an interaction of *BDNF*, *CREB1*, *GNAS*, and *POMC* genes with exposure to chronic stress or traumatic life events increase the risk of cardio-metabolic and mood disorders either simultaneously, or through mediating factors. The CRH signaling pathway is an important mediator of stress responses (124). Following an exposure to stress, the hypothalamus releases CRH, stimulating secretion of adrenocorticotrophic hormone (ACTH) from the anterior pituitary gland ACTH. This in turn stimulates the adrenal gland to produce glucocorticoids (principally cortisol). Cortisol will then act on several organs including the brain through its receptors (124). In acute conditions, the production of cortisol helps the body to fight pathogens (stress) and alleviate inflammation. However, when stressors are long lasting (chronic) they can cause cortisol receptor resistance and failure of the HPA-axis negative feedback mechanism. This increases the duration and chronicity of inflammation, and a failure to down-regulate the inflammatory response. Ultimately, failure in the HPA-axis processes may cause dysfunctions in the brain and body, causing both somatic and brain disorders. Hence, it is imperative to recognize the sources of the stress, especially chronic stress. Stress can either originate from the external environment as chronic extrinsic stress (CES) or within the internal body system as chronic intrinsic stress (CIS). Both CES and CIS can influence the CRH pathway genes mainly through gene expression and DNA methylation mechanisms (125). In relation to stress, there are two possibilities to explain mood disorders to cardio-metabolic diseases association. The first is that the human body system may consider the CMD-Rs as CIS and then dysregulate the HPA-axis through the CRH signaling pathway. Another possibility is that CES and/or CIS interact with the CRH signaling genes to cause both CMD-Rs and mood disorders. In either of the conditions, the CRH signaling genes interacts with the stressors to cause a dysfunction in the HPA-axis.

The second main canonical pathway was the adenosine monophosphate-activated protein kinase (AMPK) signaling pathway. This pathway regulates the intercellular energy balance. It inhibits or induces ATP consuming and generating pathways as needed. This pathway is especially important for nerve cells, as they need more energy with small energy reserves (126). Abnormalities in the pathway can disturb normal brain functioning. In animal studies, Zhu et al., 2014 showed chronically stressed mice developed symptoms related to mood and metabolic abnormalities, such as significant weight gain, heightened anxiety, and depressive like behavior. They also reported decreased levels of phosphorylated AMP-activated roteinkinase α (AMPKα), confirming the involvement of the AMPK pathway and its regulatory genes in metabolic disorders and depression (127). Recent studies also reported the activation of the AMPK pathway in rat hippocampus after ketamine treatment exerting rapid antidepressant effect (128). Major contributing CMMDh genes enriched in the AMPK pathway were *ADRA2A*, *ADRB1*, *LEP*, *CREB1* and *GNAS* genes. Variations in one or more of these genes can influence the activity of the AMPK pathway, subsequently impairing energy homeostasis in the brain and possibly in other cells (126). This can later cause energy shortage for the brain and somatic cells. Since brain cells are the most vulnerable units that require substantial amount of energy supply, any energy shortage would severely affect first the brain. Symptoms of mood change such as depressive behavior can be observed during this process. Moreover, AMP activation, for instance during stress, could induce insulin resistance promoting metabolic syndrome i.e. obesity, diabetes and cardiovascular diseases (129, 130). Hence, it is very likely that inappropriate activity of the AMPK pathway can imbalance the energy needs of the cells and be a cause to mood disorders and cardio-metabolic diseases.

Axonal guidance signaling was also among the top overrepresented canonical pathways. Axonal guidance signaling is related to neuronal connections formed by the extension of axons, which migrate to reach their synaptic targets. Axon guidance is an important step in neural development. It allows growing axons to stretch and reach the next target axon to form the complex neuronal networks in the brain and throughout the body. The patterns of connection between nerves depend on the regulated action of guidance cues and their neuronal receptors that are themselves encoded by axonal guidance coding genes. Activation of specific signaling pathways can promote attraction or repulsion and affect the rate of axon extension. One important observation in the axonal guidance pathway is the role of calcium and voltage-dependent calcium channels. The pathway is regulated by the entrance of calcium through the plasma membrane and release from intracellular calcium store. Calcium has been implicated in controlling axon outgrowth (131). CMMDh genes overrepresented in the axonal guidance-signaling pathway include the *BDNF*, *GNAS*, *GSK3B*, and *IGF1* genes. Mutant axonal guidance genes followed by abnormal axon guidance and connectivity could cause a disorder primarily in the brain and subsequently to the peripheral organs (132).

Other strong candidate mechanisms underlying mood disorders and cardio-metabolic diseases are the serotonin and dopamine receptors signaling pathways. The serotonin pathway is mainly regulated by serotonin and its receptors known as 5-hydroxytryptamine (5-HT) receptors. Serotonin is a monoamine neurotransmitter synthesized in the central nervous system and its signaling modulates several physiological processes including regulation of appetite, mood and sleep, body temperature and metabolism. The *SLC18A1*, *HTR1A* and *GNAS* gene were among the CMMDh genes involved in the serotonin receptor-signaling pathway. The *SLC18A1* gene encodes for the vesicular monoamine transporter that transports for monoamines. Its proper function is essential to the correct activity of the monoaminergic systems that have been implicated in several human neuropsychiatric disorders (133). The *HTR1A* gene encodes a receptor for serotonin, and it belongs to the 5-hydroxytryptamine receptor subfamily. Dysregulation of serotonergic neurotransmission has been suggested to contribute for the pathogenesis of mood disorders (60, 61) and it is implicated in the action of selective serotonin reuptake inhibitors (63-65). Moreover, animal studies have consistently demonstrated the influence of the serotonin pathway on both mood disorders and cardio-metabolic disorders. Ohta et al., 2011 have previously revealed as there is a converge in insulin and serotonin producing cells that can lead to metabolic diseases (diabetes) and mood disorders (134). The products of the insulin-producing cells (beta-islet cells) are involved to express the genes that synthesize serotonin, and serotonin also plays a role in the synthesis of insulin in the beta-islet cells (134).

The dopamine receptors pathway, centrally regulated by dopamine, also appears to underlie the relationship between mood disorders and cardio-metabolic diseases. Dopamine serves as a chemical messenger in the nervous system and its signaling plays important roles in processes: emotion, positive reinforcement, motivation, movement, and in the periphery as a modulator of renal, cardiovascular and the endocrine systems (135). The *SLC18A1* and *GNAS* genes are among the CMMDh genes that belong to this pathway. The dopamine-signaling pathway further induces the dopamine-DARPP32 Feedback in cAMP signaling. The central regulator of this pathway is the *PPP1R1B* gene that encodes a bifunctional signal transduction molecule called the dopamine and cAMP-regulated neuronal phosphoprotein (DARPP-32). Other important CMMDh genes in this pathway include *CACNA1D*, *CREB1*, and *GNAS*. The *CACNA1D* gene encodes the alpha-1D subunit of the calcium channels that mediates the entry of calcium ions into excitable cells. Calcium channel proteins are involved in a variety of calcium-dependent processes, including hormone or neurotransmitter release, and gene expression (136).

We also performed a gene network analysis of the CMMDh genes to the mood disorders and cardio-metabolic diseases. Based on the network analysis, the CMMDh genes were centrally involved in the link between mood disorders and the cardio-metabolic diseases. For instance, *ADRB1* and *ADRA2A* genes linked the four most common cardio-metabolic diseases (coronary diseases, hypertension, diabetes, obesity) with BPD and depressive disorder. The *CACNB2* and *CACNA1D* genes have shown network with coronary diseases, hypertension, diabetes, BPD and depression. Similarly, the other CMMDh genes acted as a hub between at least one of the cardio-metabolic disorders and BPD and/or depression (figure 2).

Overall, genes that encode for molecules involved in HPA-axis activity, circadian rhythm, inflammation, neurotransmission, metabolism and energy balance were found to play a central role to link mood disorders with cardio-metabolic diseases. It is also worth noting the impact of the environment, such as the CES and CIS, on the genes associated with these diseases. First, it could be that cardio-metabolic disorders and associated risk factors alter the intrinsic body environment, and this change interacts with the genes to cause mood disorders or influence treatment response in patients with mood disorders. Second, a biochemical change following the mutation of genes could result in both disease conditions simultaneously.

## IMPLICATIONS OF THE REVIEW FINDINGS

Knowledge of genes and molecular pathways that are shared between mood disorders and cardio-metabolic disorders could have several important implications for future research and clinical practice. It is expected that increasing sample size, and consequently increasing power, will identify many more of these variants in the near future. Here we identify four implications of our findings.

Firstly, the identification of shared molecular pathways implicated in disease susceptibility supports a growing evidence base for cross-diagnostic treatment paradigms. Shared molecular pathways could help explain recent findings of reduced cardiovascular mortality (137), or improved diabetic control (138), in MDD patients treated with SSRIs. Secondly, further exploration of overlapping molecular pathophysiology has the potential to unveil novel targets for drug development, and may give clues for the re-purposing of existing medications.

Thirdly, cardio-metabolic disorders are associated with an increased risk of poor response to standard treatments in mood disorders (139, 140). Genetic profiling for cardio-metabolic risk and stratified diagnosis of patients may help to classify treatment responders and treat them accordingly, thereby reducing the costs of ineffective exposure to medicines for the individuals and for the society. Early identification of at-risk individuals would also guide practitioner”s treatment recommendations, which may involve alternative somatic (e.g. electroconvulsive therapy, repetitive Transcranial Magnetic Stimulation, ketamine) or specific psychological therapies as first- or second line treatments.

Fourthly, studying the mechanisms of pleiotropic genes and shared pathways of mood disorders and somatic diseases could help untangle the clinical and genetic heterogeneity that characterizes these illnesses. It is possible that a “cardio-metabolic” endophenotype exists among mood disorders patients that may be identifiable through genetic profiling or analysis of blood protein biomarkers. Preliminary evidence for such a phenotype, approximating the concept of “atypical depression” characterized by increased appetite, weight gain, and increased need for sleep, is emerging(141, 142). Working towards personalized care that allows for precise diagnostic, treatment and prevention strategies, research could then focus on genetically stratified patient cohorts instead of the very diverse patient pool currently diagnosed with MDD or BPD. There is a growing consensus that such stratification approaches have the potential to substantially improve the quality of mental health research and mental healthcare over the coming decades (143).

Our review has limitations. Perhaps the most fundamental limitation was that almost all of the reviewed studies were performed in a univariate manner (single diseases approach). Secondly, the review included studies that reported positively associated genes, and neither negative findings nor inconsistent evidences were assessed. Thirdly, only meta-GWA studies were reviewed for the CMD-Rs. Hence, our review should be viewed as complementary to future mood disorders to cardio-metabolic diseases gene investigation, providing an initial thorough summary of potential pleiotropic genes.

## CONCLUSION

Our review revealed potential pleiotropic genes and biological pathways that are likely to be shared between mood disorders and cardio-metabolic diseases. While our review provides some insight into common mechanisms and the role of pleiotropic genes, in-depth understanding of how these genes (and possibly others) mediate the association between mood disorders and cardio-metabolic diseases requires future comprehensive cross-disorder research in large-scale genetic studies. This will enable us to better understand why patients suffer from multiple diseases at a time, and how multi-morbidities influence pharmacological treatment response to diseases.

## CONFLICT OF INTEREST

The authors declare that they have no conflict of interest.

## ACKNOWLEDGMENTS

The Adelaide Graduate Centre supported this work. Azmeraw T. Amare was a receipt of a postgraduate study scholarship, Adelaide Scholarship International (ASI), from the University of Adelaide.

